# Improving mRNA vaccine safety and efficiency with cationized lipid nanoparticle formula

**DOI:** 10.1101/2023.03.29.534838

**Authors:** Xu Peng, Guangneng Liao, Dongsheng Ren, Yucheng Zhou, Xiujin Wu, Yingxue Lei, Yan Zhang, Liang Chen, Chen He, Yaoyi Zhang, Hailin Yin, Guang Yang, Kai Xu

**Author notes:** Correspondence (K.X). These authors contributed equally to this work (Xu Peng, Guangneng Liao).

## Abstract

The widespread use of Covid-19 mRNA vaccines has highlighted the need to address rare but concerning side effects. Systemic off-target gene expression has been identified as a primary cause of acute adverse reactions and side effects associated with nucleoside-modified mRNA vaccines. In this study, we incorporated the permanent cationic lipid Dotap component into the mRNA-LNP formula associated with the FDA-approved mRNA vaccine Comirnaty to create a novel positively charged LNP carrier for mRNA vaccine delivery. Using the optimized LNP formula to prepare SARS-Cov-2 Spike mRNA vaccines for immunogenicity testing, Balb/c mice exhibited improved immunogenicity kinetics with initial antibody titers being lower but showing a continuous upward trend, ultimately reaching levels comparable to those of control mRNA vaccines 8 weeks after boost immunization. The mRNA vaccines encapsulated in the modified LNPs have demonstrated a superior safety profile in respect to systemic delivery of LNP constituents, off-target gene expression, and the systemic pro-inflammatory stimulation. Consequently, it may represent a safer alternative of conventional mRNA-LNP vaccines.

## 1. Introduction

Two Covid-19 mRNA vaccines (BNT162b2[1, 2] and mRNA-1273[3]), utilizing lipid nanoparticle (LNP) delivery technology, has been administered to billions of individuals worldwide, representing the first real-world application of prophylactic mRNA vaccines. The majority of reported adverse events following mRNA vaccination were mild to moderate in severity and typically included injection site pain, fatigue, headaches, and chills among other transient adverse events with a higher incidence than traditional vaccines[2–4]. However, rare but concerning side effects such as myocarditis and pericarditis have also been observed in young vaccine recipients[5–7]. Those troublesome occurrences may impact public acceptance of mRNA vaccines and represents a key issue that must be addressed in the development of next-generation mRNA vaccines.

In addition to inflammatory and reactogenic responses induced by vaccine components and impurities, molecular mimicry between the SARS-CoV-2 spike protein (CovS) and self-antigens, as well as immune disorders in susceptible individuals, systemic off-target gene expressions have been identified as primary causes of acute adverse reactions and side effects associated with nucleoside-modified mRNA vaccines[7, 8]. Intramuscular injection is an ideal administration route for mRNA vaccines because it ensures continuous expression of antigen protein at the injection site with affluent blood circulation[9, 10]. The protein expression kinetics are more consistent with the exponential-increment dosing for increasing antibody production principle[11, 12]. However, following intramuscular injection, a significant portion of the active ingredients may still leak into the bloodstream and choroid resulting in systemic delivery of mRNA-LNP and off-target expression of antigenic proteins[13]. A small amount of circulating Spike protein antigen was also detected in the blood of mRNA vaccine recipients with side effects tending to occur more frequently following boost immunizations. In mouse intramuscular injection experiments, tracer genes delivered by mRNA encapsulated in LNP were primarily expressed at the injection site but also appeared in tissues such as liver, lung, brain, heart, kidney, muscle, and extremities with varying intensities[12, 14]. Systemic leakage of mRNA-LNP and off-target expression of antigenic proteins can stimulate tissue damage, neurotoxicity and activate cytokines and complement which are primary causes contributing to fever, chills, and other transient adverse events. Therefore, the elimination of systemic delivery of mRNA-LNP and off-target expression of antigenic proteins represents one of the most effective means for reducing vaccine side effects.

The quality and quantity of ionizable lipids, which serve dual roles as adjuvants and gene delivery agents, are critical quality assurance for the efficacy of mRNA vaccines[15, 16]. Consequently, ionizable lipids customized for intramuscular administration of mRNA vaccines have been carefully designed and validated to minimize off-target expression of antigen genes in tissues other than muscular injection sites, thereby reducing the risk of side effects[14, 17, 18]. However, LNPs are prone to systemic gene delivery capabilities resulting in off-target expression of antigenic proteins and circulating antigenic proteins in blood. Modification by optimizing the N/P ratio and surface potential provides a feasible technical route to balance immune efficacy and localized gene expression[19, 20]. However, this approach can only reduce part of the mRNA-LNP off-target expression and has not achieved satisfactory results. It remains extremely challenging to eliminate systemic off-target gene expression of mRNA vaccines while maintaining their immunogenicity through improvements to ionizable lipids or N/P ratios.

Apolipoprotein E (ApoE) plays a major role in the clearance and hepatocellular uptake of physiological lipoproteins, also acts as an endogenous targeting ligand for LNPs, but not LNPs composed of cationic lipid[21]. Incorporating permanent cationic or anionic lipids into traditional LNPs endows nanoparticles with net positive or negative charges on their surface, altering their tissue-targeting properties for gene delivery[22, 23]. Experimental studies have confirmed that permanent cationic lipids do not possess adjuvant activity but can alter the tissue targeting of LNPs[22–26]. However, there are no reports on the use of permanent cationic lipids to reduce the systemic gene delivery capabilities of LNPs or their effect on the immunogenicity of mRNA vaccines yet.

We incorporated the permanent cationic lipid Dotap component into the mRNA-LNP formula associated with the FDA-approved mRNA vaccine Comirnaty to create a novel positively charged LNP carrier (LNP^⊕^) for mRNA vaccine delivery. With intramuscular administration as the chosen route, the formulation of LNP^⊕^ was optimized and screened using systemic off-target gene expression levels and immunogenicity efficacy as selection criteria. Using the optimized LNP^⊕^ formula to prepare CovS mRNA vaccines for immunogenicity testing, Balb/c mice in the LNP^⊕^ vaccine group exhibited ideal immunogenicity kinetics with initial antibody titers being lower but showing a continuous upward trend, ultimately reaching levels comparable to those of control mRNA vaccines 8 weeks after boost immunization. A good dosage relationship was observed between the dose of LNP^⊕^vaccine and its immunogenicity potency. Increased vaccine doses elicited higher antibody titers in mice than conventional mRNA vaccines without causing hepatotoxicity or systemic off-target gene expression.

## 2. Materials and methods

### 2.1. Lipids

Ionizable lipids: ALC-0315, SM-102, DHA-1 were purchased from Xiamen Sinopeg Biotech Co., Ltd. Cholesterol, Dotap, DSPC, Dotma, DMA-PEG2000, and Dlin-MC3-DMA were purchased from AVT (Shanghai) Pharmaceutical Tech Co., Ltd.

### 2.2. Animal

The animal experiments were conducted at the Laboratory Animal Center of Sichuan University. The maintenance and treatment of the animals fully conformed to the National Institutes of Health Guide for Animal Welfare in China. The study received approval from the Institutional Animal Care and Use Committee of Sichuan University in Sichuan, China. Female Balb/c mice aged 6-8 weeks were bred in-house and weighed between 18-20 grams at the start of the study. Prior to experimentation, the animals were acclimated for one week and then randomized into treatment and control groups.

For the collection of blood and preparation of serum, mice were anesthetized and blood was collected from the orbital sinus into a vacutainer containing either EDTA or no anticoagulant. The collected blood was then stored at 4°C for two hours before being subjected to centrifugation at 1,000 x g for ten minutes. The resulting supernatants were carefully transferred into new tubes in small aliquots and stored for long-term preservation in a −70°C freezer or for up to two months in a −20°C freezer.

### 2.3. Serum biochemical and blood analyses

Serum biochemistry analysis was conducted in Huaxi Ruipai Pet Hospital by using a veterinary clinical biochemical detector manufactured by IDXX. The blood routine testing was conducted by using VetScan HM5 (Abaxis).

### 2.4. mRNA synthesis

The CovS mRNA is identical to the BNT162b2 mRNA (Pfizer/BioNTech) in its codon region designated for a prefusion version of the SARS-Cov-2 Spike protein with G614D, K986P, and V987P substitutions. A DNA template of the CovS mRNA with a 76 nt poly-A tail was synthesized by Genscript and cloned into a plasmid DNA under a T7 expression cassette.

The production of the mRNA was carried out using an in vitro transcription (IVT) reaction system from Novoprotein with n1-methylpseudouridine (Hongene) as UTP substitute. The reaction was incubated at 37°C for six hours and treated with DNase I (Merck) prior to purification using the Monarch^®^ RNA Cleanup Kit (NEB). The RNA was then capped using Vaccinia Capping Enzyme (Novoprotein) for one hour in the vendor-recommended reaction buffer and reaction condition. The capped mRNA underwent further purification through oligo-dT chromatography (Sartorius) before being dissolved in 10 mM Citrate buffer at pH 4.0. The quality of the mRNA was analyzed through agarose gel electrophoresis and Qubit HS RNA assay. The final product was stored at −20°C.

### 2.5. LNP Preparation and Characterization

LNPs were prepared by combining a lipid mixture consisting of ionizable lipid, DSPC, cholesterol, and DMG-PEG with or without cationic lipid in ethanol with three volumes of mRNA in 10 mM of citrate buffer (pH 4.0) using a microfluidic device at a total flow rate of 5.2 mL/min. The resulting mixture was dialyzed against 1xPBS at pH 7.4 and 4°C for 16 hours to remove ethanol before being concentrated using 100-kDa Amicon filters. Particle size and uniformity were analyzed using a Particle size analyzer (Winner 802, Weina), while zeta potential was detected using Zeta Potential Analyzer (NanoBrook Omni, Brookhaven). The LNPs were stored at 4°C or −20°C for one month.

### 2.6. Bioluminescent imaging

Balb/c mice were administered with 50 μl of Fluc mRNA-LNPs via intramuscular injection in the hind leg. Following this, the animals received an intraperitoneal injection of 3 mg of D-luciferin (Beyotime). Ten minutes after fluorescein injection, mice were anesthetized using isoflurane and maintained under anesthesia through a nasal catheter. Bioluminescence was measured 13 to 15 minutes after fluorescein injection using a Xenogen IVIS-200 imaging system. The total amount of accumulated Fluc protein over time post-injection was calculated using Area under curve (AUC) analysis as described by Pardi et al.[12].

### 2.7. Enzyme-linked immunosorbent assay (ELISA)

Serum samples were analyzed using an indirect ELISA based on recombinant SARS-CoV-2 spike protein. Microtiter plates were coated with SARS-CoV-2 Spike Protein (40589-V08B1, Sino Biological) diluted in ELISA Coating Buffer. SARS-CoV/SARS-CoV-2 Spike Chimeric Mab (40150-D002, Sino Biological) was serially diluted as antibody references. Diluted serum samples or antibody references (100 μl) were pipetted into the Spike-coated plates and incubated at 37°C for 1 hour. Rabbit Anti-Mouse IgG H&L (HRP) (ab6728, Abcam) was then added and incubated for an additional hour. Unbound material was washed off with 1xPBS between each step and 3,5,3’5’-tetramethylbenzidine (TMB) (Abcam) was used as a chromogenic substrate to detect antibody responses. The corresponding change in color was measured at 450 nm using an Infinite F50 microplate reader (Tecan). The optical density (OD) values and the concentration of the antibody references were subsequently used to calculate the relative antibody titer in samples.

### 2.8. The TNF-α and IFN-γ ELISA

The determination of serum TNF-α and IFN-γ concentration was performed using Mouse IFN gamma Uncoated ELISA and Mouse TNF alpha Uncoated ELISA (Invitrogen). The protocols recommended by the vendor were strictly adhered to.

### 2.9. Surrogate ELISA for RBD-ACE2 competitive neutralizing antibodies

Microtiter plates were coated with SARS-CoV-2 Spike RBD-mFc Recombinant Protein (40592-V05H, Sino Biological) diluted in ELISA Coating Buffer. The His-ACE2 protein (10108-H08H, Sino Biological) was diluted in 0.5% non-fat milk solution and used as the competitor protein. SARS-CoV/SARS-CoV-2 Spike Chimeric Mab was serially diluted as antibody binding references. Serum samples or antibody binding references (100 μl) were mixed with 60 μl of competitive ACE2 and then pipetted into the RBD-coated plates at 37°C for 1 hour. Unbound material was washed off with 1xPBS. HRP-labeled Anti-His tag Antibody (105327-MM02T-H, Sino Biological) was then incubated at 37°C for an additional hour. TMB was used as a chromogenic substrate and OD was measured at 450 nm using an Infinite F50 microplate reader.

Samples containing SARS-CoV-2 neutralizing antibodies inhibit the protein–protein reaction between His-ACE2 and the RBD competitively. The average OD value of the sample is subtracted from the average OD value of the blank control to calculate the calibration value of the sample. The OD value of control is designated as B0, and the OD value of the well with inhibitor added is designated as B. The percentage of B/B0 is referred to as the binding rate, and the corresponding inhibitor concentration with calculated binding rate as 50% is determined to be IC50 value.

### 2.10 Statistic Analyses

The statistical analysis of experimental data and independent unpaired *t*-tests were performed utilizing IBM SPSS Statistics v20.

## 3. Results

### 3.1. LNP^⊕^s reduced systemic off-target gene expression

By testing the effect of intramuscular injection of LNP^⊕^ preparations on the delivery and expression patterns of Firefly luciferase (Fluc) mRNA, we identified LNP^⊕^25, LNP^⊕^46 and LNP^⊕^74 formulations with comparable expression levels at the injection site to traditional LNP preparations but with very low or no expression in liver, lung and brain and other visceral tissues (Figure 1). The lipid composition and physicochemical properties of the nanoparticles are listed (Figure 1a). Control LNP was formulated with an mRNA vaccine using ionizable lipid ALC-0315.

**Figure 1.**
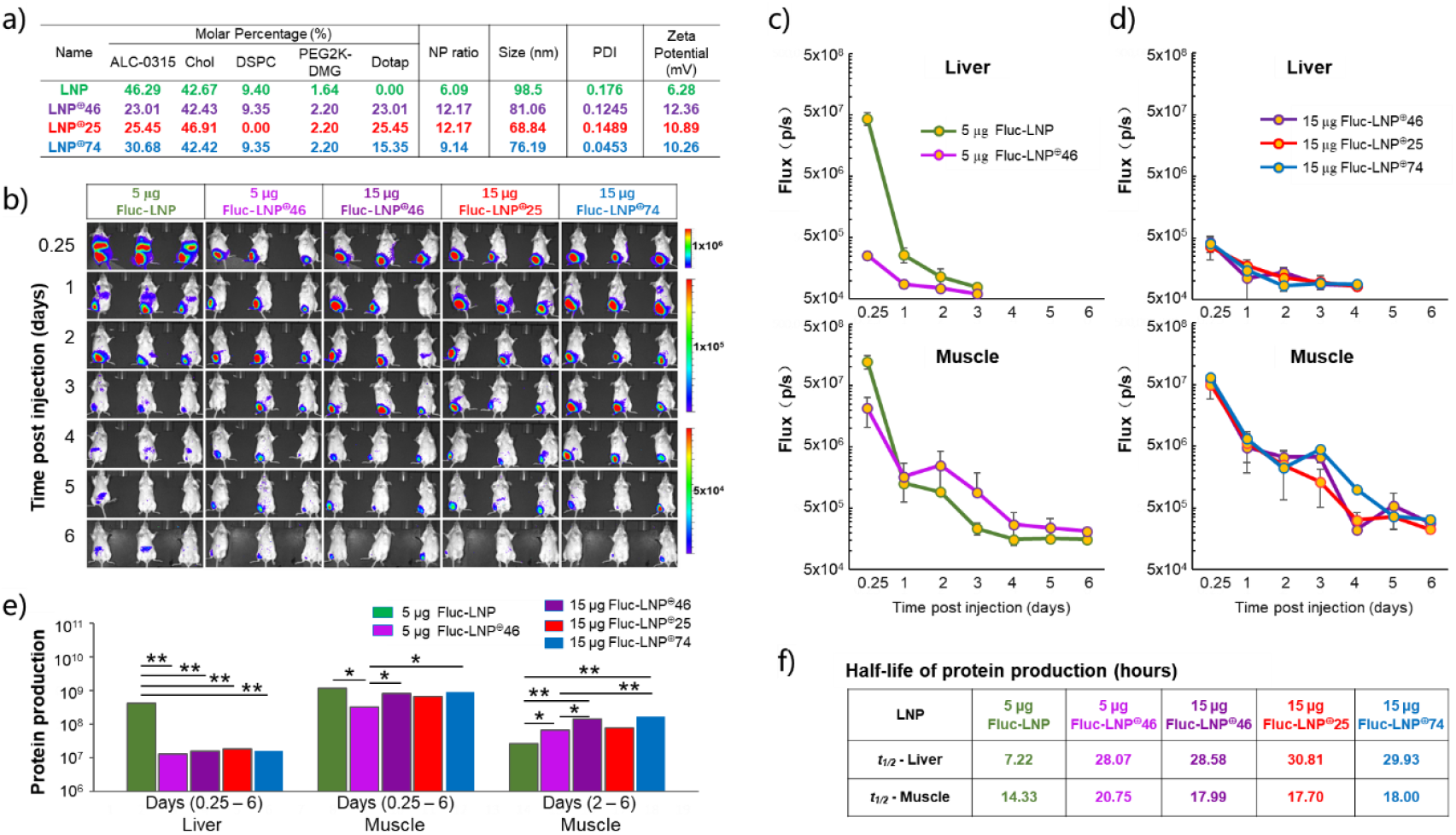
Expression kinetics of FLuc mRNA in LNPs with and without cationic Dotap lipid. (a) Composition and particle size of LNPs. (b) Bioluminescent images of mice acquired via IVIS imaging at 0.25, 1 to 6 days after intramuscular injection (n = 3 mice per group). (c) and (d) Flux counts in Liver and Muscle measured at 0.25, 1 to 6 days after intramuscular injection of 5 μg and 15 μg Fluc mRNA, respectively. (e) Total amount of protein over different time frames calculated using the area under the curve (AUC). Statistical significance between cohorts determined by independent unpaired *t*-tests (*p <0.05, **p < 0.01). Error bars represent mean ± SEM. (f) Half-life of protein over different time frames in Liver and Muscle, respectively, in relevant timelines.

The permanent cationic lipid Dotap and the ionizable lipid exhibit similar physical and chemical properties[27]. Consequently, LNP^⊕^ with analogous properties was assembled utilizing established LNP microfluidic techniques. Measured in 20% Glucose solution, the surface potential of LNP^⊕^s was significantly elevated in comparison to that of LNP (Figure 1a). The increased surface charge enhances the affinity between nucleic acid molecules and positively charged lipids for improved structural integrity of the LNP^⊕^. We found that mRNA-LNP^⊕^ partially disintegrates in Triton solutions with concentrations below 7.5% and completely dissolves in 10% Triton solutions. Since encapsulation efficiency measurements for LNP were usually conducted in a 1% Triton solution, so mRNA-LNP^⊕^ dissociation rates at varying Triton concentrations were evaluated and a detection method was developed for LNP^⊕^ encapsulation efficiency by using Qubit HS RNA Kit with 10% Triton as the lysate (Supplementary Table 1-3).

Live IVIS imaging revealed significant alterations in the distribution of gene expression delivered by mRNA-LNP^⊕^ following intramuscular injection in Balb/c mice. At a dose of 5 μg mRNA, Fluc gene expression was primarily observed at the intramuscular injection site and liver in the LNP control group mice (Figure 1b). Expression decreased to background levels within 48 hours, consistent with previously reported gene expression results by LNPs[12]. In contrast, the LNP^⊕^46 administration group exhibited Fluc gene expression at the intramuscular injection site for up to 9 days post-administration, with no detectable expression in liver tissue.

Figure 1c illustrates the Fluc expression intensity curve in mouse liver and intramuscular injection site at a dose of 5 μg mRNA between 6 hours and 6 days post-administration. Protein expression was quantified using the fluorescein intensity of AUC. Within 24 hours, protein expression in LNP group mouse liver tissue was 31.81 times higher than that of the LNP^⊕^46 administration group. However, protein production at the intramuscular injection site was 2.55 times higher in the LNP^⊕^46 group compared to controls between days 2 and 6, indicating a significant shift in gene expression kinetics (Figure 1e). Supplementary Figure 2 provides details on the visceral distribution of Fluc gene expression and demonstrates that DSPC played a minor role in promoting systemic gene expression.

In comparison to the low-dose group, high-dose administration (15 μg mRNA) of LNP^⊕^ exhibited the similar gene expression kinetics (Figure 1d). However, protein expression at the intramuscular injection site increased significantly and exhibited a positive correlation with mRNA usage and protein production (Figure 1e). The enhanced and sustained protein expression facilitated by LNP^⊕^ fits better with the principle of exponential-increment dosing for higher antibody production.

### 3.2. LNP^⊕^s formed from various ionizable lipids had similar expression kinetics

The surface potential of Dotap-LNP^⊕^s undergoes a transformation to exhibit a weak positive charge. This change repels LNP^⊕^s from conjugation with ApoE and other lipid carriers in the blood, diminishing their systemic transportation. It has been postulated that incorporating permanent cationic lipids into LNPs with varying ionizable lipid compositions might achieve the same effect and could represent a general approach. The gene delivery efficacy of Dotap and Dotma is inferior to that of ionizable lipids[26, 28, 29]. When LNPs containing only Dotap or Dotma were administered to mice via intramuscular injection, only low-level tracer protein signals were detected at the injection site (Figure 2a). The incorporation of Dotap or Dotma into LNPs composed of different ionizable lipids such as MC3, DHA-1, L319, SM-102 was analyzed using live IVIS imaging following intramuscular injection. Results indicated that incorporation of permanent cationic lipids significantly reduced protein expression in mouse liver tissue delivered by LNP; this phenomenon was observed universally (Figure 2b-f).

**Figure 2.**
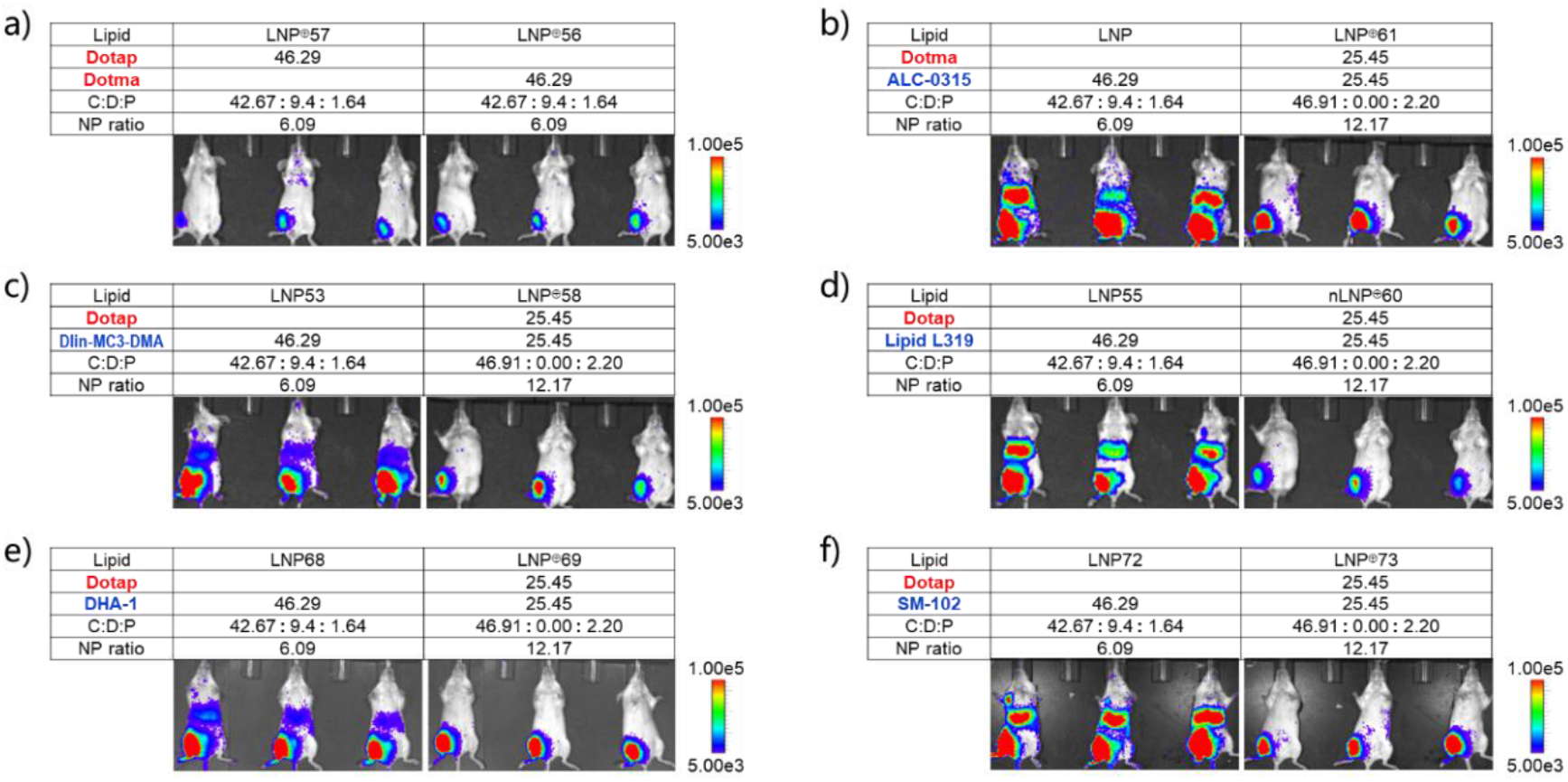
Expression kinetics of LNPs formulated with different ionizable lipids, with and without permanent cationic lipids. LNPs containing either Dotap or Dotma alone (a) and LNPs formulated using one of the five lipidoids: ALC-0315 (b), Dlin-MC3-DMA (c), Lipid L319 (d), DHA-1 (e), and SM-102 (f) with lipid compositions listed in molar ratio. mRNA-LNPs administered intramuscularly to Balb/c mice at a dose of 0.375 mg/kg (n = 3 mice per group). Six hours later, mice were injected with D-luciferin for live IVIS imaging analysis. Lower panels exhibit the representative IVIS images of Fluc expression.

### 3.3. LNP^⊕^ formula induced Lymphocyte Homing

Intramuscular injection of LNP^⊕^ significantly reduced the expression level of delivered genes in liver tissue. To investigate its effect on blood immune cells and liver tissue, 20 μg of CovS mRNA encapsulated either in LNP or LNP^⊕^ were administered to mice via intramuscular injection. Blood samples were collected from the mice and laboratory tests were performed on blood cell counts and liver injury-related biochemical markers. The results showed significant changes in four blood parameters: WBC count, lymphocyte count, ALT, and AST (Figure 3a-d). Changes in serum levels of TNF-α and IFN-γ were also detected (Figure 3e,f). in addition, no changes in body weight or behavioral abnormalities were observed in any of the mice vaccinated with LNP or LNP^⊕^ during this study.

**Figure 3.**
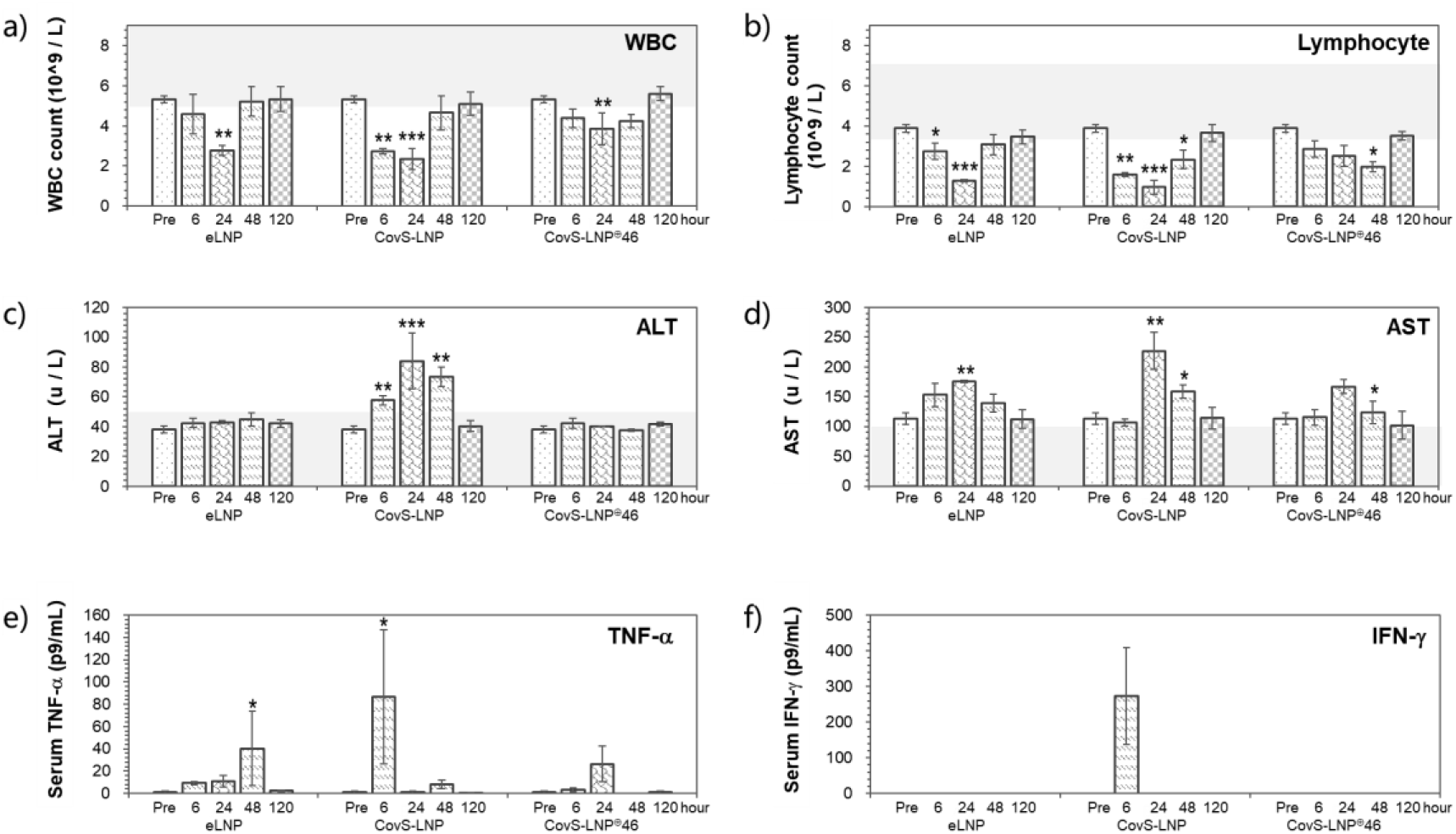
Hepatotoxic and immunological biochemical biomarkers in Balb/c mice significantly altered following intramuscular administration of 20 μg CovS mRNA encapsulated in LNPs. LNP without mRNA (eLNP) was used as a negative control. (a) White blood cell count (WBC), (b) lymphocyte count, (c) Alanine Aminotransferase (ALT), (d) Aspartate Aminotransferase (AST), (e) TNF-α, and (f) IFN-γ were measured at pre-treatment and 6, 24, 48, and 120 hours post-administration with mean ± SEM calculated. Physiological ranges of parameters of Balb/c mice illustrated as gray boxes. Statistical significance between each cohort and healthy control determined by independent unpaired *t*-tests (*p < 0.05, **p < 0.01, ***p < 0.001).

Clinical trials have indicated that lymphocyte homing was induced by the administration of the BNT162b2 vaccine. Blood samples from mRNA vaccine recipients showed a transient decrease in lymphocyte counts that returned to normal levels within one week[2]. A complete blood count analysis on mouse blood samples indicated a significant reduction in blood lymphocyte count in the CovS-LNP group. The number of lymphocytes decreased significantly six hours after LNP administration, reaching its lowest in 24 hours. However, it returned to normal levels in six days (Figure 3b). The CovS-LNP^⊕^46 group showed a gradual decline in lymphocyte counts and reached the lowest in two days and returned to normal levels in six days, demonstrating slower reduction kinetics of lymphocytes in circulation. The eLNP group showed a reduction pattern similar to that of the CovS-LNP positive controls, but to a lesser degree. This elucidated that the lymphocyte homing was induced largely by the LNP shell rather than the antigen expressed. In all groups, the changes of white blood cell counts agreed with the lymphocyte reduction, indicating that lymphocytes were the major target of homing induced by mRNA-LNP. LNP^⊕^ induced lymphocyte homing but in a milder way.

### 3.4. Blood biochemical analysis for Hepatotoxicity and pro-inflammatory cytokines

In the CovS-LNP control group, blood ALT levels increased significantly six hours after intramuscular injection and peaked after 24 hours, indicating the occurrence of liver injury (Figure 3c). The AST also increased significantly at the same time (Figure 3d). In the eLNP group, AST increased slightly but ALT remained unchanged. Hemolysis and inflammation at the injection site may have caused the slight increase in AST rather than liver injury. No clear signs of liver injury were observed in the eLNP and CovS-LNP^⊕^46 groups, indicating that expression of S protein antigen in liver tissue was responsible for the liver injury in mice.

Results indicated that mice in the CovS-LNP group exhibited a rapid and transient increase in serum TNF-α and serum IFN-γ levels, peaking at six hours and returning to baseline after 24 hours (Figure 3e,f). in a previous study, mice vaccinated with 10 μg mRNA-1273 showed increased TNF-α and IFN-γ levels in muscle 24 hours later; however, only serum IFN-γ levels were significantly elevated[30]. BNT162b2 vaccination similarly yielded elevated muscle TNF-α expression without notable changes in serum levels. Our study indicated that transient increases in serum levels of TNF-α and IFN-γ at the six-hour mark coincided with the peak of systemic off-target gene expression resulting from CovS-LNP administration, while no such elevations were observed in the CovS-LNP^⊕^ group. It is thus inferred that this surge in systemic off-target gene expression might account for the brief elevation in serum TNF-α and IFN-γ levels.

### 3.5. LNP^⊕^ formula improved CovS mRNA immunogenicity

The gene distribution and expression kinetics resulting from intramuscular injection of LNP^⊕^ are distinct from those of traditional LNP and are more consistent with the exponential-incremental dosing for higher immunogenicity principle. We used CovS mRNA to evaluate the efficiency of LNP^⊕^ vaccine-stimulated antibody production on Balb/c mice. The particle size and encapsulation efficiency of the LNP^⊕^ in use was listed in Supplementary Table 4.

An immunization experiment comprising two immunizations was conducted and is illustrated in Figure 4a. Mice in the CovS-LNP group were observed to generate a modest quantity of S protein-specific IgG antibodies three weeks following initial immunization. A significant rise in antibody titers was observed one week after boost immunization, peaking before gradually declining (Figure 4b). In contrast, mice in the CovS-LNP^⊕^74 group demonstrated a small but significant number of antibodies one week after boost immunization, representing 6.5% of the titers of the control CovS-LNP group during this same period. However, an upward trend in immune reactivity was noticed, and reaching the same level as that of LNP controls 8 weeks after boost immunization, corresponding to the sustained gene expression dynamics and slow homing trend of blood lymphocytes associated with LNP^⊕^. Notably, increasing the mRNA dose to 15 μg led to significantly enhanced immunogenicity of the LNP^⊕^74 vaccine, resulting in the titers three times higher than that of the control group even 8 weeks after boost immunization. A clear correlation between antibody titer and mRNA dosage was observed.

**Figure 4.**
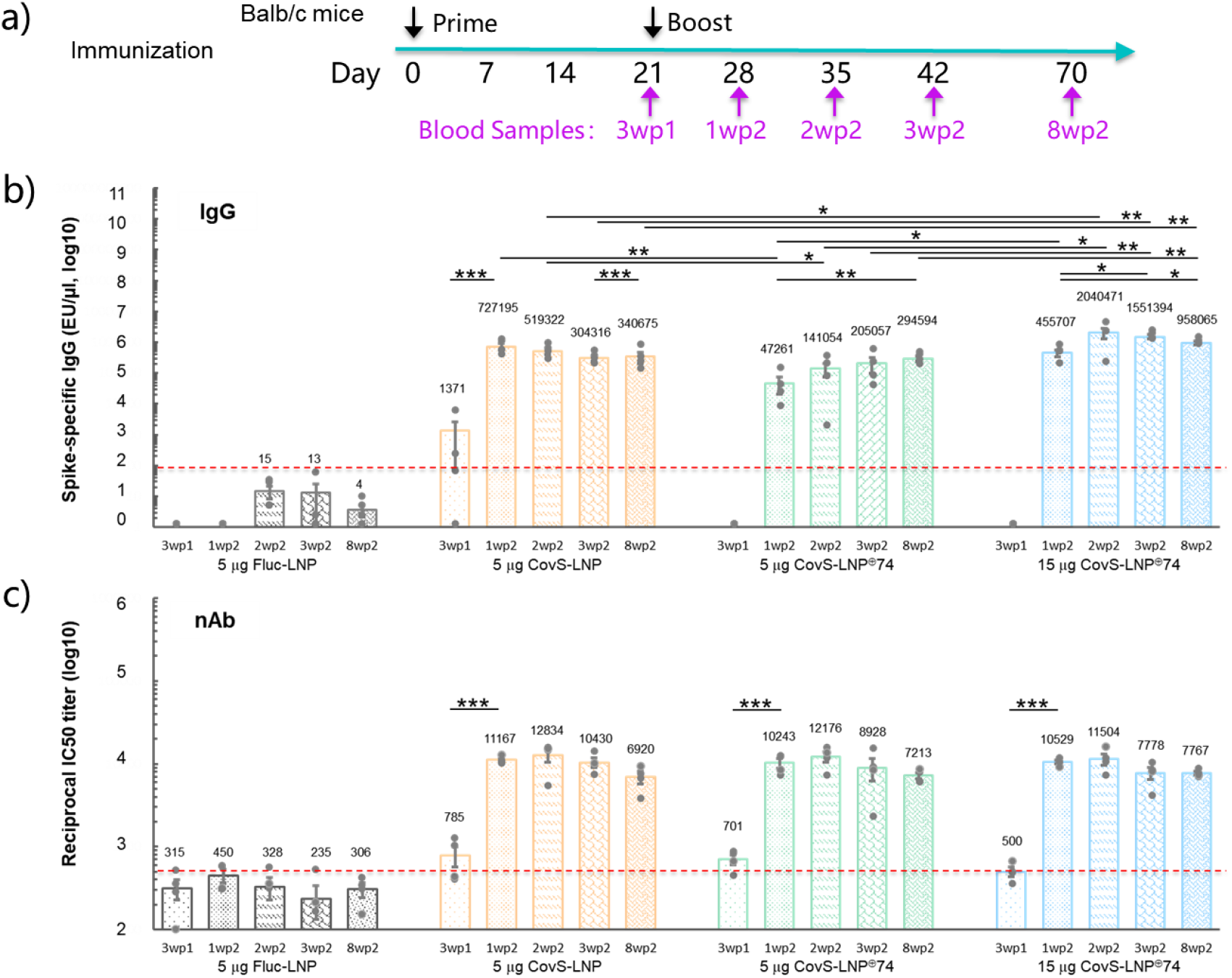
Immune response in CovS mRNA encapsulated in LPN or LNP^⊕^74 vaccinated mice. Balb/c mice administered intramuscularly with different amounts of mRNA-LNPs in 30 μl of 1xPBS, as well as an equivalent dose 21 days later as a booster (n = 4 mice per group). (a) Schematic schedule of immunization and sample collection. (b) IgG titer determined using a Spike-protein specific binding antibody ELISA assay. (c) Reciprocal IC50 titer assessed by an RBD-ACE2 competitive binding ELISA. Data shown as mean ± SD. Each dot represents an individual animal. Dashed line indicates detection limit of assay. Significance was calculated using independent unpaired *t*-tests (*p < 0.05, **p < 0.01, ***p < 0.001). Difference between negative controls and corresponding experimental groups not indicated.

To further reinforce these findings, a surrogate ELISA assay for RBD-ACE2 competitive neutralizing antibody was performed on serum samples from immunized mice as previously suggested[31]. Elevated levels of neutralizing antibody titers were observed one week after boost immunization in mice receiving various doses of LNP^⊕^74- or LNP-encapsulated mRNA vaccines. Titers peaked two weeks after boost immunization, and gradually declined (Figure 4c). Surprisingly, no significant differences were observed in the neutralizing antibody titers between mice treated with two doses of LNP^⊕^74. To ascertain the consistency of the results, another set of serum samples was tested at different dilutions. Higher IgG binding antibody titers were achieved in line with the CovS mRNA dosage; however, the absence of a dose-corresponding increase in neutralizing antibody titers persisted, although it remained at its peak 8 weeks after boost immunization. One possible explanation for this observation is that the CovS mRNA dosage used in the experimental settings was excessive for Balb/c mice. Mice in the negative control group consistently produced little to no neutralizing antibodies, well below the assay LOD.

With respect to systemic delivery, off-target gene expression, and the systemic stimulation of pro-inflammatory factors by LNP platform, LNP^⊕^ technology has produced a superior safety profile. A higher dose of CovS mRNA encapsulated in LNP^⊕^ elicited greater IgG antibody titers than its LNP counterpart, with fewer concerns regarding side effects in treated mice. Consequently, mRNA-LNP^⊕^ vaccines may represent a safer alternative that provides stronger and long-lasting immune protection than conventional mRNA-LNP vaccines.

## 4. Discussion

Clinical studies have revealed a weak correlation between reactogenicity and immunogenicity of mRNA vaccines, as well as a lack of direct link between T cell responses and adverse events of mRNA vaccines[32]. The common adverse reactions of nucleoside-modified mRNA vaccines are independent of immunogenic efficacy, which offers feasibility for the development of safer, more effective next-generation mRNA-LNP vaccines[33].

At physiological pH, LNPs rely on a neutral surface to achieve systemic gene delivery through ApoE-based blood transport carriers for efficient liver-specific gene delivery. Upon intramuscular injection, LNPs may leak into the bloodstream and lymphocytic ducts through the penetration of active LNP components from mRNA vaccines, potentially causing adverse events and side effects. The incorporation of permanent cationic lipids alters the surface charge properties of LNP^⊕^s, reducing their systemic transportation capacity and promoting mRNA retention and expression at the injection site while maintaining particle stability. However, permanent cationic lipids lack adjuvant functionality and cannot replace ionizable lipids in stimulating immune responses. To ensure immunostimulatory potency, a sufficient amount of ionizable lipids, which serve as adjuvants in LNPs, must be included in the LNP formula aimed at vaccination.

LNP^⊕^ formula possesses adjuvant functionality and is evidenced in its ability to induce comparable lymphocyte homing. At equivalent dose levels, the CovS mRNA encapsulated in LNP^⊕^74 elicited a gradual increase in S protein-specific binding antibody titers, eventually reaching levels comparable to those of the LNP control vaccine 8 weeks after boost immunization. However, neutralizing antibody levels were comparable to those of the LNP control vaccine from the outset and slightly higher in higher dose of CovS-LNP^⊕^74 group in 8 weeks. Using the amount of tracer protein as a reference, the total antigenic protein produced by mice receiving 15 μg of CovS-LNP^⊕^74 should be lower than that of the 5 μg CovS-LNP vaccine group. However, the antibody titers produced in mice of the former group were three times higher. This phenomenon may be attributed to alterations in the kinetics of gene expression delivered by LNP^⊕^74, which enhanced the incremental mode and duration of antigenic protein accumulation, resulting in a gradual increase in antibody titers and a more balanced immunogenic response. Simultaneously, the slow production of antigenic proteins may help to avoid an instantaneous impact on the immune system, reducing the immune burden caused by innate immune responses and decreasing the risk of cytokine storms and acute adverse events.

The Spike protein binds to the ACE2 receptor, potentially causing detrimental effects on cells that express the receptor. In this study, we administered CovS mRNA via intramuscular injection using LNPs and found that off-target expression of the CovS antigen protein in liver tissue was the primary cause of liver injury in mice. Additionally, systemic off-target gene expression resulted in a transient increase in serum levels of TNF-α and IFN-γ. Systemic inflammation mediated by inflammatory cytokines or cardiac protein mimicry has been linked to myocarditis[34, 35] and a higher incidence of adverse events associated with the mRNA-1273 vaccine compared to the Comirnaty vaccine[30]. The use of LNP^⊕^ significantly diminished systemic off-target expression of antigenic proteins, thereby preventing liver damage and cardiac inflammation, and reducing the production of systemic pro-inflammatory factors. Although comprehensive safety profiling of LNP^⊕^ formula has not yet been completed, our data suggests that the risk of adverse reactions from LNP^⊕^ vaccines is significantly lower than that of traditional LNP vaccines.

In conclusion, the incorporation of Dotap into traditional LNPs containing ionizable lipids can effectively reduce systemic gene delivery and off-target gene expression, improving gene expression kinetics. This may provide a general approach for the development of mRNA vaccines by altering gene-targeted delivery through modification of surface potential. This model can be extended and applied to various permanent cationic lipid molecules, such as Dotma, DC-Chol, DOSPA or their derivatives, as a general strategy for developing intramuscularly injected mRNA vaccines. However, further research is needed to confirm the safety and efficacy of LNP^⊕^ in human subjects and to fully explore its long-term effects on immune protection and potential side effects.

## 5. Conclusion

This study presents a novel approach to improving the safety and efficacy of mRNA vaccines by incorporating the permanent cationic lipid into the mRNA-LNP formula to create a positively charged LNP carrier for mRNA vaccine delivery. Results demonstrate that the novel LNPs have a superior safety profile compared to traditional LNP vaccines, with reduced systemic off-target gene expression and systemic inflammatory stimulation.

## Author Contributions

All authors confirmed they have contributed to the intellectual content of this paper with final approval of the publication. K.X., D.R., and G.Y conceived the concept and prepared the manuscript. Y.C.Z., X.W., C.H., H.Y., and K.X. designed the experiments and analyzed data. Y.L., Y.Y.Z., and Y.Z developed the LNP Encapsulation Efficiency assay and carried out the production of mRNAs. X.P., G.L., Z.F., and L.C design and carried out the animal experiments.

## Acknowledgements

The authors thanked Jian Yang and Lindan Diao for their technical assistances.

