## Supplementary File for "Improving mRNA vaccine safety and efficiency with cationized lipid nanoparticle formula"

This PDF file includes:

1. The quantification of RNA and DNA in various Triton X solutions
2. Method for Determination of Encapsulation Efficiency of LNP and LNP$^{\oplus}$
3. Supplementary Figure 1. LNP$^{\oplus}$ Stability
4. Supplementary Figure 2. DSPC influences Systemic gene expression of LNP$^{\oplus}$
5. Properties of mRNA-LNP$^{\oplus}$ used in immunization

### 1. The quantification of RNA and DNA in various Triton X solutions

The Qubit HS RNA assay (Q32852, Invitrogen) is a reliable and straightforward method for the quantitative detection of RNA. However, the impact of high concentrations of detergent on the accuracy of detection has not been reported. The Qubit Fluorometer 2.0 (ThermoFisher) was utilized to quantitatively detect a custom standard RNA solution (with a quantitative detection value of 270 ng/ml in 1xTE solution) in various concentrations of Triton solutions. The detection results are presented in Supplementary Table 1.

Supplementary Table 1. RNA quantification in various Triton X solutions

| Triton X-100 (v/v%) | Test1 | Test2 | Test3 | Average | SD |
|---|---|---|---|---|---|
| 0.0% | 269 | 266 | 266 | 267.0 | - |
| 0.1% | 272 | 270 | 270 | 270.7 | 1.39% |
| 0.05% | 271 | 270 | 271 | 270.7 | 1.39% |
| 0.01% | 268 | 269 | 267 | 268.0 | 0.37% |

| | | | | | |
|---|---|---|---|---|---|
| 0.005% | 274 | 275 | 272 | 273.7 | 2.51% |
| 0.002% | 274 | 274 | 273 | 273.7 | 2.51% |
| 0.001% | 275 | 276 | 272 | 274.3 | 2.73% |

The findings indicate that Triton X-100 with a final concentration ranging from 0.001% to 0.1% had no significant effect on the RNA quantification results obtained using the Qubit HS RNA Kit, with a coefficient of variation below 3%. Owing to the dilution factor, the concentration of Triton that can be used in RNA detection samples ranges from 0% to 20%. These results were also replicated and confirmed in the quantitative detection of DNA using the Qubit HS dsDNA Kit (Q32851, Invitrogen).

### 2. Method for Determination of Encapsulation Efficiency of LNP and LNP$^{\oplus}$

The encapsulation efficiency of LNP and LNP$^{\oplus}$ is determined using the Qubit Fluorometer 2.0 in conjunction with the Qubit RNA HS Assay Kit and the Qubit dsDNA HS Assay Kit. The RNA content before and after dissolution of LNPs or LNP$^{\oplus}$s in 10% Triton X-100 solutions is quantitatively detected. The encapsulation efficiency is then calculated based on the RNA contents detected before and after LNP or LNP$^{\oplus}$ dissolution. The procedure is as follows:

a) Determination of RNA concentration (A): An equal volume of 20% Triton X-100 solution is added to the LNP or LNP$^{\oplus}$ sample to be tested, mixed well and centrifuged, and left at room temperature in the dark for 5 minutes. The sample is diluted 200 times for detection, and the total RNA concentration in the lysed sample is obtained as RNA concentration (A).

b) Determination of RNA concentration (B): Without adding Triton detergent, the content of unbound RNA in the LNP or LNP$^{\oplus}$ sample is detected as the RNA concentration (B) .

c) Calculation formula: Encapsulation Efficiency (%) = ((A - B) / A) × 100

LNP is completely dissolved in a low-concentration Triton solution, so the detection of the encapsulation efficiency of LNP is generally carried out in a Triton solution with a concentration below 2%. LNP$^{\oplus}$ has increased stability to non-ionic detergents, and the RNA content and lipid components can only be completely separated in 10% Triton solution. At different detergent concentrations, we tested and analyzed the encapsulation efficiency of LNP and LNP$^{\oplus}$ with an RNA content of 0.1 μg/μl. The nucleic acid content test results are shown in Supplementary Table 2.

Supplementary Table 2: The RNA content encapsulated in LNPs detected in different concentrations of Triton solution

| | | LNP | LNP$^{\oplus}$28 | LNP$^{\oplus}$25 | LNP$^{\oplus}$31 | LNP$^{\oplus}$32 | LNP$^{\oplus}$33 | LNP$^{\oplus}$34 | LNP$^{\oplus}$35 | LNP$^{\oplus}$37 |
|---|---|---|---|---|---|---|---|---|---|---|
| Cationic Lipid | Dotap | | 25.59 | 25.45 | 25.31 | 25.24 | 24.29 | 23.33 | 22.29 | 32.30 |
| Ionizable Lipid | ALC-0315 | 46.29 | 25.59 | 25.45 | 25.31 | 25.24 | 24.29 | 23.33 | 22.29 | 21.53 |
| Help Lipid | Cholesterol | 42.67 | 47.17 | 46.91 | 46.65 | 46.53 | 48.95 | 51.33 | 53.50 | 39.70 |
| | DSPC | 9.4 | - | - | - | - | - | - | - | 4.37 |

| | | | | | | | | | | |
|---|---|---|---|---|---|---|---|---|---|---|
| | PEG-DMG | 1.64 | 1.66 | 2.22 | 2.73 | 3.00 | 2.11 | 2.02 | 1.93 | 2.09 |
| The RNA concentration detected from LNPs prepared with the given formula in different Triton solution | | | | | | | | | | |
| 1% Triton | A | 0.100 | 0.029 | 0.033 | 0.029 | 0.041 | 0.051 | 0.031 | 0.031 | 0.044 |
| 5% Triton | A | 0.103 | 0.032 | 0.054 | 0.042 | 0.044 | 0.059 | 0.061 | 0.049 | 0.032 |
| 7.5% Triton | A | 0.098 | 0.081 | 0.085 | 0.073 | 0.88 | 0.86 | 0.81 | 0.082 | 0.077 |
| The RNA concentration detected from LNPs prepared with the given formula in a 10% or 0% Triton solution | | | | | | | | | | |
| 10% Triton | A | 0.112 | 0.101 | 0.102 | 0.116 | 0.102 | 0.104 | 0.113 | 0.101 | 0.112 |
| 0% Triton | B | 0.016 | 0.001 | 0.001 | 0.003 | 0.001 | 0.001 | 0.002 | 0.001 | 0.001 |
| EE (%) | | 85.451 | 99.335 | 98.965 | 97.446 | 98.58 | 99.067 | 97.879 | 99.278 | 98.635 |

In Supplementary Table 2, the concentration unit of lipid components is expressed in mmol%. The N/P ratio of LNP is 6, while the N/P ratio of all LNP$^{\oplus}$ is 12. B and A represent the detection of RNA concentration values determined before and after LNP$^{\oplus}$ dissociation by detergent (unit: μg/μl).

The results indicate that a 1% Triton solution can completely dissolve traditional LNP but is unable to fully dissolve LNP$^{\oplus}$. The degree of LNP$^{\oplus}$ dissociation increases with the increase in Triton concentration. A 10% Triton solution completely decomposes LNP$^{\oplus}$ prepared by using different formulations, and the total nucleic acid content is consistent with the theoretical total nucleic acid in the sample.

Comparative experiments with plasmid DNA-LNP$^{\oplus}$ yielded similar results. As shown in Supplementary Table 3, a 10% Triton solution does not affect the quantitative detection of plasmid DNA using the Qubit HS dsDNA Kit detection method.

Supplementary Table 3. The detected value of DNA content in DNA-LNP and DNA-LNP$^{\oplus}$ in various concentrations of Triton

| | | LNP | LNP$^{\oplus}$25 | | |
|---|---|---|---|---|---|
| DNA (μg) | | 0.15 | 0.30 | 0.15 | 0.075 |
| 1% Triton | A | 0.111 | 0.0044 | 0.0023 | 0.0082 |
| 10% Triton | A | 0.118 | 0.315 | 0.153 | 0.072 |
| 0% Triton | B | 0.00073 | 0.00113 | 0.00080 | 0.003 |
| EE (%) | | 99.39 | 99.64 | 99.48 | 99.28 |

B and A in Supplementary Table represent the DNA concentration values determined before and after LNP$^{\oplus}$ (or LNP) dissociation (unit: μg/μl).

**3. Supplementary Figure 1. LNP Stability**

The results of the LNP stability test showed that LNP and LNP$^{\oplus}$ stored at 2°C - 8°C for 2 weeks had no significant effect on gene delivery and expression.

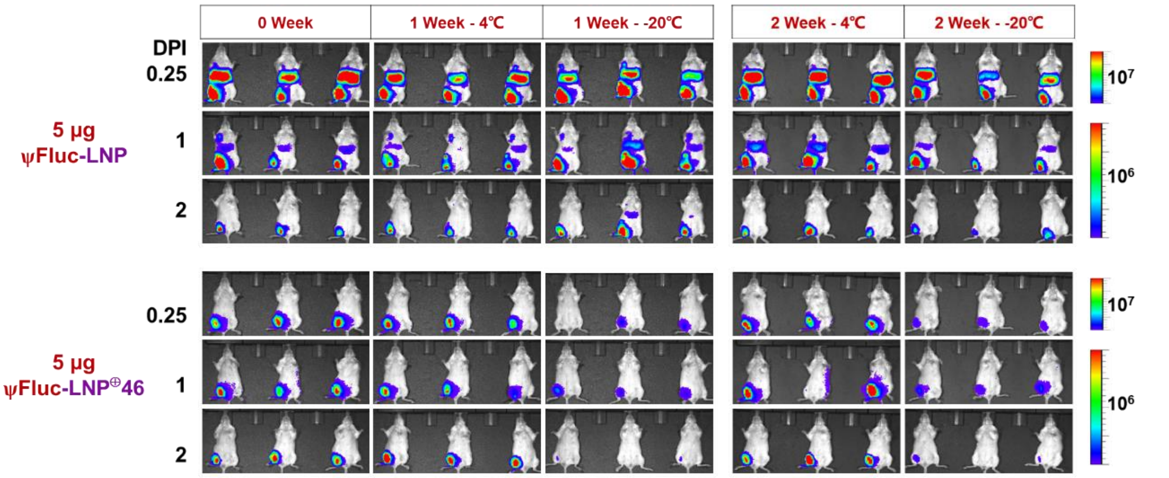


Supplementary Figure 1. Expression intensity of Fluc-LNP and Fluc-LNP⊕46 stored at 4°C or -20°C over 0, 1, and 2 week periods (n = 3 mice per group). Mice were injected with D-luciferin for live IVIS imaging analysis at 6, 24, and 48 hours post-intramuscular injection.

### 4. Supplementary Figure 2. DSPC influences Systemic gene expression of LNP⊕

It has been observed that the content of DSPC exerts a minor impact on systemic gene expression. Fluc mRNA was encapsulated using LNP⊕s formulated with varying DSPC contents and subsequently tested on Balb/c mice. The component contents for LNP⊕25, LNP⊕45, and LNP⊕46 were identical, with the exception of DSPC content, which was 0.00%, 4.91%, and 9.35%, respectively.

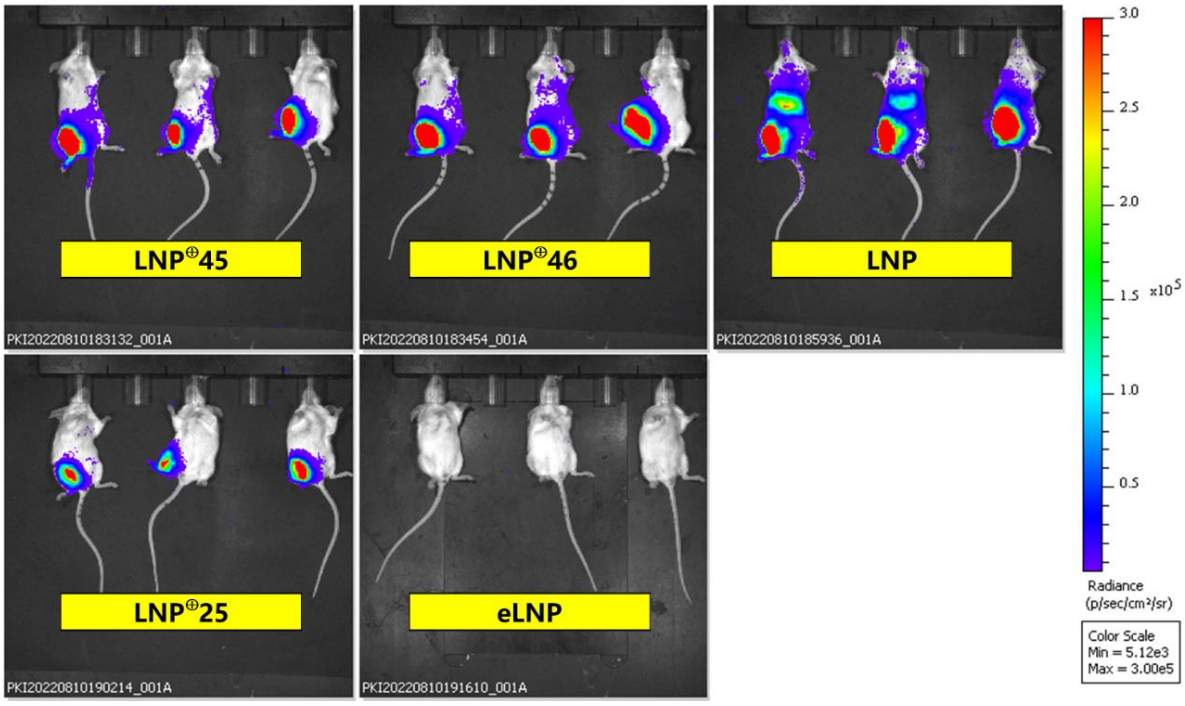


Supplementary Figure 2. Distribution of Fluc expression six hours after the intramuscular injection of 5 μg of Fluc mRNA encapsulated in LNP and LNP⊕s.

### 5. Properties of mRNA-LNP$^{\oplus}$ used in immunization

Supplementary Table 4. Size, PDI and Encapsulation efficacy of LNP$^{\oplus}$s used in mouse immunization experiment

| LNP | Formula | mRNA Class | Size (nm) | PDI | EE % |
|---|---|---|---|---|---|
| Fluc-LNP | LNP | Fluc mRNA | 90.83 | 0.1061 | 90.23 |
| CovS-LNP | LNP | CovS mRNA | 74.14 | 0.1028 | 96.61 |
| CovS-LNP$^{\oplus}$46 | LNP$^{\oplus}$46 | CovS mRNA | 77.61 | 0.1124 | 96.31 |
| CovS-LNP$^{\oplus}$25 | LNP$^{\oplus}$25 | CovS mRNA | 73.79 | 0.1398 | 97.46 |
| CovS-LNP$^{\oplus}$74 | LNP$^{\oplus}$74 | CovS mRNA | 81.31 | 0.1162 | 98.71 |
